# Evaluation of the Biological Effects and Specificity of the Novel Dihydroceramide Desaturase 1 inhibitor PR280 and its analogues

**DOI:** 10.64898/2026.11.25.768879

**Authors:** Deanna M. Peperno, Hector R. Romero Garcia, Charly N. Campanella, Nihal Medatwal, Pablo Rivero, Yolanda Díaz, M. Isabel Matheu, Christopher J. Clarke

## Abstract

Emerging evidence suggests that dihydroceramide desaturase (DES1) plays key roles in many cellular biologies and pathologies, underscoring a need for DES1 inhibitors, both as research tools and potential therapeutic agents. In this study, we have evaluated the recently reported novel DES1 inhibitor PR280 and its newly synthesized analog PR280b. We find that while both PR280 and PR280b are potent in vitro and in situ DES1 inhibitors, they induced broader changes in cellular sphingolipids – most notably increased ceramide levels – that go beyond DES1 inhibition and had modest effects on DES1-driven biologies including anchorage-independent survival, apoptosis, and autophagy. Overall, although this limits the direct utility of PR280 as a specific DES1 inhibitor, this highlights the potential of this scaffold for further inhibitor development aimed at maintaining the DES1 inhibitory activity while reducing off-target effects.

---

Dihydroceramide desaturase (DES) occupies the last key step of the *de novo* sphingolipid (SL) synthesis pathway, catalyzing the introduction of the C4-C5 *trans* double bond and converting dihydroceramide (dhCer) to ceramide (Cer) (1). At present, two mammalian DES isoforms have been identified and cloned. Of these, DES1 (gene name: DEGS1) shows ubiquitous expression and is the primary activity in many tissues (2). In contrast, DES2 (gene name: DEGS2) is most highly expressed in skin and exhibits both desaturase and 4-hydroxylase activity, being able to produce both dhCer and phytoceramide, respectively (3, 4). Compared to other SL enzymes, the DES isoforms have been less well studied, primarily as dhCer was thought to be inactive due to early studies with short-chain dhCer analogues that showed no biological activity (5, 6). The discovery that *N*-(4-hydroxyphenyl)retinamide (4HPR)-induced autophagy and cell death was associated with accumulation of dhCer, rather than Cer (7, 8) led to renewed interest in dhCer. Since then, many studies have overturned this early perception, establishing dhCer as a versatile signaling molecule with uniquely complex and sometimes contradictory cellular functions. At the cellular level, loss of DES1 has been associated with a number of anti-tumor biologies including activation of apoptosis (7, 9), growth arrest (10, 11), autophagy (12, 13), and ER stress pathways (14, 15). DES1 can also impact other functions including cell migration, cytoskeleton modification, insulin response, and impaired endomembrane trafficking (13). Pathologically, DES1 has been implicated in breast, colorectal, lung, and brain cancers (9, 14–16) with a recent study suggested DES1 is important for metastasis of colon cancer (16). We also identified DES1 as a key target of oncogenic HER2 and showed that loss of DES1 leads to a decrease in anchorage-independent-survival (AIS) and anchorage-independent growth (AIG) (14) – two key metastasis-enabling biologies. Outside of cancer, considerable evidence both from human and animal studies has implicated Cer accumulation with lipotoxic cardiometabolic disease and liver fibrosis (17–21) and shown that targeting DES1 was effective at restoring insulin sensitivity and overcoming these adverse effects (19–21). Taken together, this growing literature demonstrates the high therapeutic potential for specifically targeting DES1.

The emerging roles of DES1 in various cellular biologies and disease states emphasizes the growing potential for DES1 inhibitors as therapeutic agents. Moreover, as DES1 knockout mice are embryonic lethal (22), DES1 inhibitors would have added value as research tools, particularly for studying *in vivo* functions. Nonetheless, the availability of potent and specific DES1 inhibitors remains limited. Follow on studies based on the connection of dhCer with 4HPR-induced cell death established that 4HPR - a synthetic vitamin A analogue with well-established chemotherapeutic properties - was a direct *in vitro* inhibitor of DES1 (23) at lower concentrations than 4HPR is typically used. Nonetheless, 4HPR is well known to affect many other targets such as retinoid receptors (24) mTOR (25), JNK (26) and FAK (27), as well as having poor bioavailability (28) and so its utility is limited. Certainly, for cell studies, it should be used at low doses (around 2μM) and complemented by genetic manipulation with siRNA, shRNA, and CRISPR knockout. Following on from 4HPR, GT11 was developed as the first specific DES1 inhibitor and is a C8-*N*-acylated cyclopropenyl Cer that inhibits DES1 in rat liver microsomes with an IC_50_ of 20 µM (29) but has limited chemical and biological stability which also limits its use (30). It was also reported to have off-target effects albeit at higher doses (31). Like GT11, XM462 is a Cer analogue that contains a sulfur-containing group in place of Cer’s C4–C5 double bond, a modification inspired by sulfur-containing fatty acids that inhibit other desaturases (30, 32). XM462 provides mixed type inhibition and was reported to have IC_50_ of 0.43 μM in Jurkat cells (30, 32). However, cell studies suggested that it did not seem to appreciably alter sphingolipids in the presence of serum and accumulated both dhCer and Cer levels in serum-free conditions, suggesting off-target effects (30, 32). Additionally, structure-function studies led to identification of the compound CIN038 as a DES inhibitor (33) but although subsequent studies showed it led to increased dhCer levels, it did not lead to decreased Cer levels (34) as is typically seen with DES1 loss. More recently, the cyclopropenone-containing Cer analogue PR280 was identified as a novel DES1 inhibitor with an IC_50_ of 700 nM in T98 cell lysates, which exceeded that of GT11 and XM462 as reference inhibitors (35). Structural modeling and docking supported key noncovalent interactions within the DES1 active site, providing a potential basis for its inhibitory activity. However, the cellular consequences of Pr280 treatment and its effects on DES1-associated biologies remain unclear.

Building on these findings, the present study investigates PR280 and its newly synthesized analogs, assessing their inhibitory activity against DES. Additionally, the effects of the most active compounds on cellular SL levels and DES1-regulated biologies including AIS, AIG, apoptosis and autophagy were also assessed.

## MATERIALS AND METHODS

### Reagents

Standard laboratory reagents were from Millipore Sigma or Fisher Scientific. Anti-DES1/MLD antibody (ab167169) was from Abcam and anti-actin was from Millipore Sigma.

### Cell lines and culture

JIMT-1 cells (DSMZ ACC-589) cells were obtained from DSMZ-German Collection of Microorganisms and Cell Cultures GmbH, respectively, and authenticated by short tandem repeat (STR) profiling. JIMT-1 cells were cultured in Dulbecco’s Modified Eagle Medium (DMEM; #11965092, Gibco/Thermo Fisher Scientific); SKBR3 cells were cultured in Dulbecco’s Modified Eagle Medium Ham’s F12 (DMEM/F-12; #11320033, Gibco/ Thermo Fisher Scientific) respectively, supplemented with 10% heat-inactivated fetal bovine serum (FBS; #26400044, Thermo Fisher Scientific). Cell lines were maintained at 37°C in a humidified atmosphere containing 5% CO₂ and passaged at approximately 85% confluence using 0.05% Trypsin-EDTA (#25300054, Gibco/Thermo Fisher Scientific) following washing with Dulbecco’s phosphate-buffered saline (DPBS; #14190250, Thermo Fisher Scientific). Cells were routinely tested for mycoplasma contamination using a luminescence-based assay. All experiments were performed using cells between passages 3 and 25.

### Synthesis of PR280 and structurally related analogues

Cyclopropenone ceramide analogues PR280a and PR280b were synthesized as described for the reference compound PR280 (36). Specifically, aminodiol **1** (**Scheme 1**) was subjected to *N*-acylation with hexanoic and decanoic acid, using EDC and HOBt as coupling agents, affording the desired products in 73% and 66% yield, respectively. Full details for synthesis, purification of products, and spectroscopical data are provided in Supplemental Information.

**Scheme 1.**
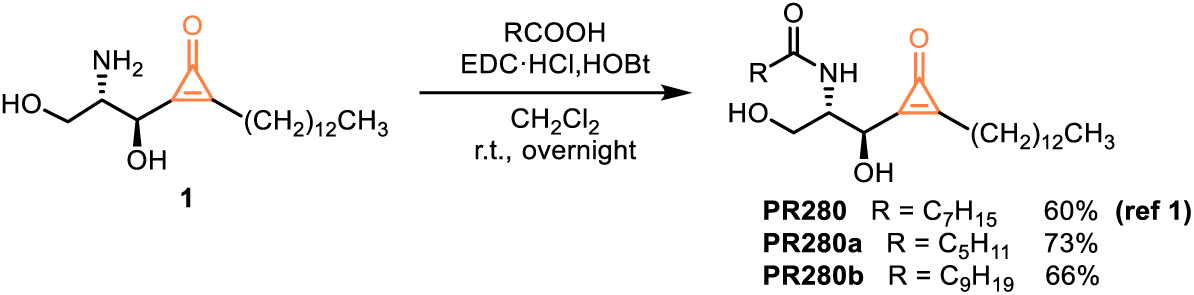
Synthesis of cyclopropenone ceramide analogues **PR280**, **PR280a, PR280b** via *N*-acylation of aminodiol **1**.

### siRNA transfection

Cells were plated into 100 mm dishes at a density of 800K cells per dish. siRNA complexes were prepared using Lipofectamine RNAiMAX in OptiMEM according to the manufacturer’s protocol. Briefly, siRNA was diluted in OptiMEM and mixed with an equal volume of RNAiMAX diluted in OptiMEM. The complexes were incubated for 15 minutes at room temperature prior to being added dropwise to the dishes. All Star siRNA (Qiagen) was used as a negative control, and all siRNAs were used at a final concentration of 20nM. Following transfection, cells were incubated for 48 hours before media change (1-2h) and treatment as shown.

### Generation of DES1 knockout JIMT1 cell lines

Lentiviral particles were generated by seeding 2.5 × 10⁶ HEK293T cells in 100 mm culture dishes. Twenty-four hours later, cells were co-transfected with 1.5 µg target plasmid, 1.5 µg pCMV-VSV-G, and 1.5 µg pCMV-dR8.2 DVPR using Lipofectamine 2000 (3:1 ratio) according to manufacturer instructions. After 24h, media was replaced and 24h later, 2ml of media was added. At 72h post-transfection, virus-containing supernatants were collected and filtered (0.45 µm PVDF; #SLHVR33RS, Millipore Sigma), aliquoted, and stored at −80C. For infection, JIMT1 cells (150K) were seeded in 6-well plates. The following day, cells were treated with 1ml viral supernatant combined with 1 ml media containing polybrene (8 µg/ml final concentration). The next day, media was changed and 48h post-infection, cells were passaged into media containing puromycin (1 µg/ml) for 7-8 days. Polyclonal stable cell populations were used for all downstream experiments.

### Analysis of proteins by SDS-PAGE and Western Blot

Cellular proteins were extracted using RIPA buffer (50 mM Tris, 150 mM NaCl, 1% Triton X-100, 0.1% SDS, protease inhibitor, phosphatase inhibitor). Samples were sonicated for 10 seconds to disrupt cells, and protein concentration was determined by Bradford assay using BioRad reagent (#50000006). Equal volumes of lysates were mixed with 2× Laemmli buffer (#1610737, BioRad) containing 5% β-mercaptoethanol (#M6250, Millipore Sigma), heated to 100 °C for 10 minutes, and stored at −20°C until subsequent analysis. For SDS-PAGE, 7.5–20 µg of protein was loaded onto 4–20% acrylamide gels with 5 µL molecular weight markers between sample sets. Proteins were separated at room temperature at a constant current of 50 mA for approximately 1.5 hours, then transferred onto nitrocellulose membranes by wet transfer in transfer buffer (20% methanol, 25 mM Tris, 192 mM glycine, pH 8.3) at 100 V for 1 hour at 4 °C. Membranes were blocked in 5% milk for 1 hour, rinsed with 0.1% PBS-T, and incubated overnight at 4 °C with primary antibodies diluted in 2% bovine serum albumin in 0.1% PBS-T. The following day, membranes were washed (3 × 5 minutes, 0.1% PBS-T) and incubated with HRP-conjugated secondary antibody (1:5000 in 2% bovine serum albumin in 0.1% PBS-T) for 1.5 hours at room temperature. After additional washes (3 × 5 minutes, 0.1% PBS-T), membranes were developed using enhanced chemiluminescence (ECL, #32109, ThermoFisher Scientific) for 1-10 minutes and exposed to X-ray film.

### Analysis of dihydroceramide desaturase activity by *in situ* assay

Measurement of *in situ* DES activity was performed using cell-permeable C12-dhCCPS substrate as described previously (15, 37). Briefly, cells were plated and treated according to the experimental conditions being evaluated and for the final hour of treatment, C12-dhCCPS (final concentration: 500nM) was added. To extract lipids, cells were washed with ice-cold PBS, scraped into 2 ml of Cell Extraction Mix A (2:3 70% isopropanol:ethyl acetate), and transferred into 15 ml Falcon tubes. To prepare lipids for mass spectrometry analysis, 10 µl of C12-dihydroceramide (#860635, Avanti Research) was added to each sample as internal standard, followed by an additional 2 mL of Cell Extraction Mix A. Samples were vortexed thoroughly, centrifuged (3000 rpm, 5 min, 4 °C) and supernatants transferred to glass tubes. After drying under nitrogen, dried extracts were reconstituted in 150 µl of Mobile Phase B (methanol with 1% formic acid and 1 mM ammonium formate). Levels of C12-dhCCPS (substrate) and C12-CCPS (product) were separated and analyzed using tandem HPLC lipid chromatography-mass spectrometry as previously described (3). In situ DES activity was calculated as the ratio of product to substrate.

### Analysis of dihydroceramide desaturase activity by *in vitro* assay

Measurement of *in vitro* DES activity was performed as described previously (14, 36). Briefly, JIMT-1 cells were harvested in phosphate-buffered saline (PBS) and pelleted by centrifugation (5 min, 3500rpm, 4 °C). Cell pellets were resuspended in lysis buffer (5 mM HEPES pH 7.4, 50 mM sucrose), lysed by sonication on ice (3 x 10s pulses), and unbroken cells and debris removed by centrifugation (250 xg, 5 min, 4 °C). Protein concentration was determined by Bradford assay. For each reaction, glass test tubes containing 50 μg of protein were adjusted to 100 μL with lysis buffer, with blank control reactions prepared using 100 μl of lysis buffer without protein. A reaction mastermix was containing 5mM HEPES pH 7.4, 50mM Sucrose, 2μM C12-Pyr dhCer substrate and 2mM NADH (freshly prepared each time and dissolved in lysis buffer). To initiate reactions, 100ul of mastermix was added to blank and lysate samples (final reaction volume of 200μl) and samples were incubated at 37 °C for 20 minutes. The enzymatic reaction was terminated with 2 ml cell extraction mix A (see above) and extracted for mass spec analysis according to the same method as for in situ assays above.

### Analysis of cellular SLs

After treatment, cells were washed once with cold 1X PBS and scraped in 2ml of extraction buffer A (2:3 70% Isopropanol: Ethyl acetate). Extract was transferred to 15 ml tubes and stored at −80 °C before submission to the Lipidomics core. To further extract for analysis and quantification, samples were bought up to room temperature and 50 μl of appropriate internal standards were added. An additional 2ml of extraction buffer A was added, samples were vortexed, and centrifuged (5 min, 3500 rpm). The resultant organic phase was split into glass tubes (3 ml for lipid analysis, 1 ml for phosphate analysis) and dried under liquid nitrogen. The 3 ml dried extract was reconstituted in 150 μl of methanol and submitted to the Lipidomics Core for analysis by tandem liquid chromatography/mass spectrometry (LC/MS) at the Stony Brook Lipidomics Core as described previously (14, 37). The dried 1 ml extracted was processed for analysis of total lipid phosphate levels, analyzed as described previously (14, 37) for subsequent normalization of total lipid levels.

### Analysis of Golgi SL flux by NBD assay

This assay was performed as described previously (39). Briefly, NBD-C_6_-Ceramide (Cayman Chemical 62527) was first conjugated to fatty acid-free BSA in PBS. For this, 20 μl of 4.5 mM NBD-C_6_-Cer in ethanol was added to 1 ml of 0.34mg/ml fatty acid-free BSA in PBS, vortexed, and stored at −20 °C. For experiments, cells were plated and treated as desired with 1μM NBD-C_6_-Cer/BSA complex added to cell culture medium for the last 1h of incubation. Cells were washed with PBS and scraped in 2ml of lipid extraction solvent A (2:3 70% isopropanol: ethyl acetate), transferred to a 15ml conical tube, and stored at −80C. For further extraction, samples were thawed at room temperature before addition of 2ml of extraction solvent A. After vortexing and centrifugation (3000 rpm, 5 min), organic layers (approx. 4 ml) were transferred to glass tubes. A one ml aliquot was removed for inorganic phosphate measurements while the remaining 3ml extract was dried down under nitrogen gas and resuspended in 100μl of mass spectrometry grade methanol for subsequent HPLC analysis.

### Analysis of cell death by trypan blue exclusion

Cells in monolayer (Att) were harvested by trypsinization, reactions stopped by addition of equal volume of culture media, and pelleted by centrifugation (1200 rpm, 5 min). Cells cultured in ECM-detached conditions (Susp) were harvested from trays and pelleted by centrifugation (1200 rpm, 5 min). Cell pellets were vigorously resuspended in 2ml trypsin-EDTA and left at room temperature for 1-2 mins to dissociate cell clumps and produce a single cell suspension. Reactions were stopped with 2ml of complete media, and dissociated cells were pelleted by centrifugation. At this point, cell pellets from monolayer and suspension were vigorously resuspended in 1ml of growth media. 20 μl of cell suspension was combined with 20 μl 0.4% trypan blue solution (#15250-061, GIBCO, Life Technologies) for 30-45s and 10 μl of the mix was used for cell counting by hemocytometer. Cells that showed no presence of dye (trypan blue negative) were counted as living. Cells with showing presence of dye in the interior (trypan blue positive) were counted as dead. Data are expressed as % trypan blue positive cells of total cells counted.

### Colony formation in soft agar

This was performed as described previously (14). Briefly, a lower layer of 0.8% agar was created by mixing 2x media and supplements with an equal volume of sterile 1.6% agar in dH_2_O. Once lower layers had solidified, cells were plated in an upper layer of 0.4% agar (30K for SKBR3; 10K for 10A-NeuT). For this, appropriate cell numbers were added to 2x media and mixed with an equal volume of 0.8% agar. Once solidified, trays were incubated for 2-3 weeks at 37 °C as detailed. 200 μl of normal growth media was added to the top of each well every 3-4 days to ensure they did not dry out. Once experiments had completed, cells were stained overnight with nitrotetrazolium blue chloride (#N6876, Millipore Sigma). Images of 5-6 fields per condition were taken at 4x and 10x magnification with an EVOS microscope for each replicate, and number of visible colonies were counted. For inhibitor experiments, the appropriate 2x volume of inhibitor was added to cells and 2x media prior to mixing with 0.8% agar.

### Statistical Analysis

Data were graphed and analyzed using Graphpad Prism. Analysis of two means was conducted with Student’s t-test; analysis of more than two means by one-way ANOVA with appropriate post-test; and comparison of two variables by two-way ANOVA with appropriate post-test. For all statistical tests, a *p*-value of less than 0.05 was considered as threshold sufficient to reject the null hypothesis.

## RESULTS

### PR280 analogues inhibit *in situ* and *in vitro* DES activity

Prior research assessed the potential for PR280 and other analogues with a rigid scaffold as inhibitors of DES1, combining previous structure-activity studies of other DES1 inhibitors (GT11, XM462) with AlphaFold2-assisted structural insights (32). As PR280 showed promise as a DES inhibitor in T98G cells (IC_50_ = 0.7 μM), here we expanded investigations into two additional PR280 analogues – namely PR280-a and PR280-b – that were synthesized as described in detail in Supplemental Methods. To begin to assess these analogues relative to parental PR280, we utilized a well characterized *in situ* DES activity assay that we have utilized before (14, 33) and chose JIMT1 breast cancer cells as a cell system, primarily as DES1 expression is over 100 times higher than DES2 in this cell line and over 99% of the *in situ* activity detected by our assay is due to DES1 (data not shown). Thus, we can say that observed effects of compounds on in situ activity are likely a consequence of DES1 inhibition. Initial activity assays with the three compounds suggested that PR280 and PR280-b had stronger effects on DES activity than PR280-a (data not shown). Given this, a full dose response (0, 1, 2, 5, 10 μM) of PR280 and PR280-b on *in situ* DES activity was performed (**Fig. 1A**). For PR280, results showed approx 97.3% inhibition at 1 μM treatment with slight – but not statistically significant – increases in inhibition at higher doses. Similar effects were seen with PR280-b with 96% inhibition at 1 μM and slightly stronger effects at 2, 5 and 10 μM. Analysis of DES1 protein levels showed that compounds did not decrease DES1 levels and, if anything, led to an increase in DES1 protein over vehicle control. This is consistent with post-translational inhibition rather than induced degradation of the protein (**Fig. 1B**). To build on these data, we wanted to see how acutely each analogue could inhibit DES and so performed a time course. Here, cells were treated for 2, 4, and 6h with either Cer analogue (or vehicle control) prior to assessment of *in situ* DES activity (**Fig. 1C**).

**Figure 1.**
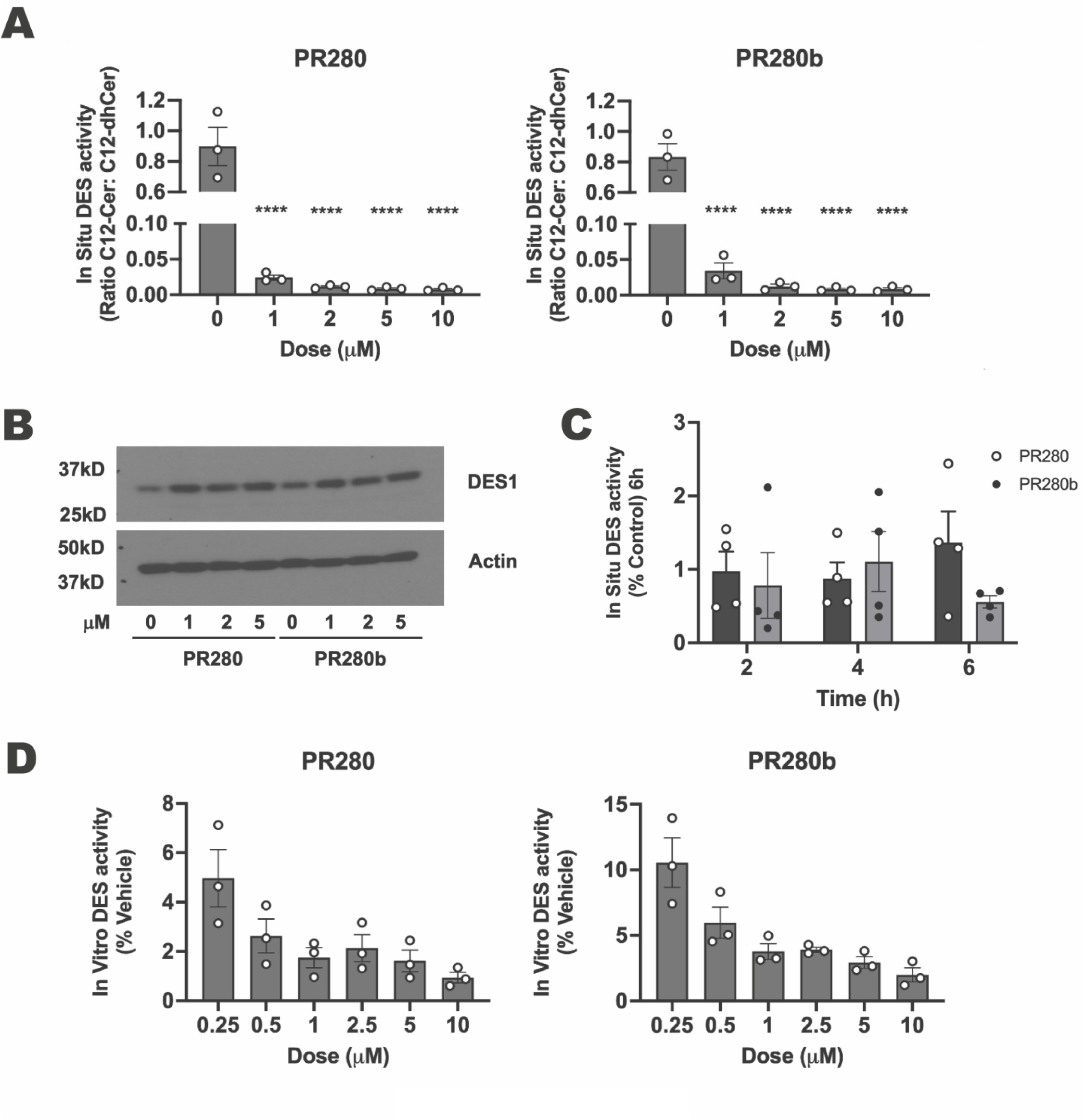
PR280 analogues inhibit in situ and in vitro DES activity at low to submicromolar concentrations. (A) JIMT1 cells were treated with PR280 or PR280b for 24 h with the indicated doses and *in situ* DES activity was analyzed. Data are expressed as mean ± SEM *in situ* activity (Ratio product: substrate) (**** p<0.0001, one-way ANOVA with Dunnet’s post test; n = 3) (B) JIMT1 cells were treated with PR280 or PR280b for 24 h with the indicated doses. Protein was extracted and analyzed by immunoblot as shown. Representative of at least 3 biological replicates (C) JIMT1 cells were treated with 2μM PR280 or PR280b for the times shown and *in situ* DES activity was analyzed. Data are expressed as mean ± SEM % vehicle control activity at 6h (n = 4) (D) JIMT1 lysate was incubated with PR280 or PR280b for 10 min with the indicated doses prior to analysis of in vitro DES activity. Data are expressed as mean ± SEM % vehicle control activity (n = 4)

As *in situ* results suggested a greater *in situ* DES1 inhibition with PR280 than what was observed previously in T98G cells (35), we assessed effects on *in vitro* DES1 activity utilizing JIMT1 cell lysate as protein source and testing the range of 0.25μM to 10μM (**Fig. 1D**). Strikingly, for PR280, as little as 250nM of inhibitor led to a substantial inhibition of DES activity (>96%) and this was observed for all doses tested. Stronger *in vitro* effects were also observed with PR280-b although as with *in situ* activity, the effects were not as robust as PR280 with approximately 93% inhibition observed at 250nM. Taken together, these data support PR280 and PR280b as effective inhibitors of DES1.

### Effects of PR280 analogues on cellular sphingolipid levels

As we observed stronger DES1 inhibition of both analogues at lower concentrations than prior studies (32), it was important to determine their effects on endogenous SL levels. To this end, we treated JIMT1 cells with 2 μM of each compound for 24 h, utilizing a DES1 Crispr knockout cell line for comparison for direct comparison (**Fig. 2**). Results indicated that both inhibitors led to a robust increase in dhCer levels comparable to that seen in DES1 KO cells and consistent with DES1 inhibition (**Fig. 2A**). Both inhibitors and DES1 KO led to reduced hexosylceramide (HexCer) (**Fig. 2F**) and sphingomyelin (SM) levels (**Fig. 2G**), also consistent with DES1 inhibition as this is immediately upstream of these reactions in the de novo SL pathway. Similar reductions of sphingosine (**Fig. 2C**), and S1P levels (**Fig. 2D**) were also seen with inhibitors and DES1 KO although it should be noted that both inhibitors had modestly higher effects on sphingosine (Sph). However, while DES1 KO caused a reduction in total Cer levels, as would be anticipated from a loss of DES1 activity, both PR280 and PR280-b caused an increase in Cer levels (**Fig. 2B**). Similarly, DES1 KO also caused a significant increase in dihydrosphingosine (dhSph) levels, presumably due to a back-up of de novo SL synthesis and as seen previously (14) yet treatment with PR280 and PR280-b led to decreased dhSph levels (**Fig. 2E**).

**Figure 2.**
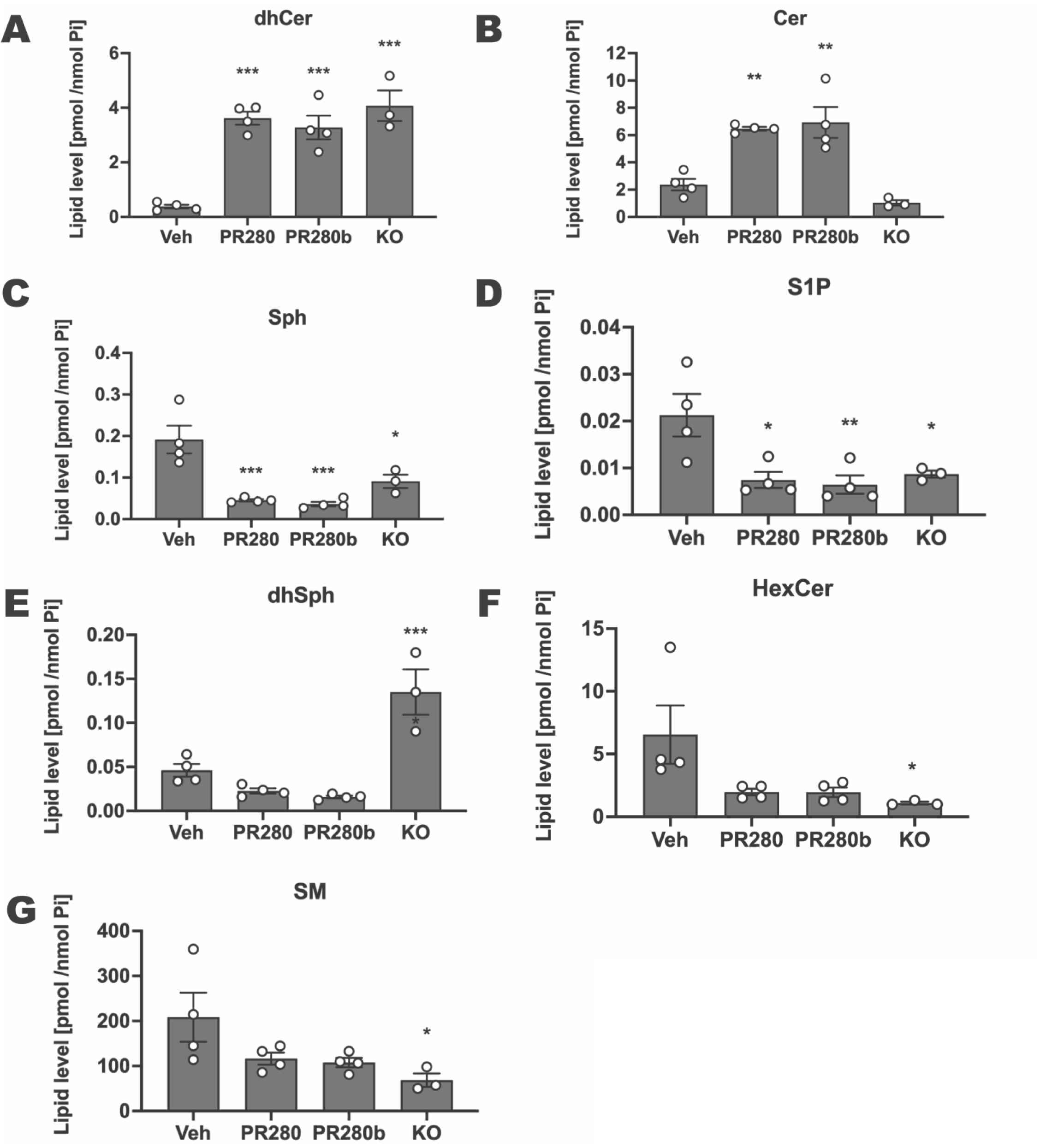
Effects of PR280 and PR280b on sphingolipid profiles compared to DES1 knockout cells. JIMT1-Vec and JIMT1-DES1 KO cells were cultured for 24h in the presence of 5 μM PR280, PR280b or vehicle control as shown. Lipids were extracted and analyzed for the lipid classes shown. Data are expressed as mean ± SEM of lipid level normalized to total inorganic lipid phosphate (n = 4; * p< 0.05; ** p < 0.01, *** p < 0.001 vs vehicle; one-way ANOVA with Dunnet’s post-test).

To extend these data and determine if the observed inhibitor-driven changes in lipids are tied to effects on DES1, we utilized siRNA to knockdown DES1 with AllStar siRNA as negative control. After 48 h, both control and knockdown cells were treated with 2 μM PR280 or PR280b for 24 h (**Fig. 3**). Here, effects of DES1 knockdown with siRNA were similar to DES1 KO cells leading to significant accumulation of the upstream lipids dhCer and dhSph, with significant reductions in downstream lipid classes of Cer and HexCer. In contrast to DES1 KO cells, effects of DES1 KD on Sph and S1P were more modest but did decrease, albeit not significantly. In si-AS control treated cells, both analogs had similar effects to the previous experiments, causing increases in dhCer and Cer levels, but decreases in Sph, dhSph, S1P, and HexCer levels. Of note, the increases in dhCer levels for both compounds were comparable to vehicle-treated si-D1 cells. However, effects in si-D1 treated cells appeared to be more complex. For dhCer, levels were increased further in analogue-treated si-D1 cells compared to si-AS cells. In contrast, Cer levels were reduced in analogue-treated si-D1 cells compared to inhibitor-treated si-AS cells – but notably were still elevated by both compounds compared to Veh control in si-D1 cells. In contrast, for dhSph, Sph, and S1P, there was no difference in analogue effects between si-AS and si-D1 cells (with strong reduction seen in both cases) while HexCer levels were modestly reduced further in analogue-treated si-D1 cells compared to si-AS cells. Taken together, these results suggest wider effects of both PR280 and PR280b on the SL network that are independent of DES1 – most notably effects on cellular Cer levels.

**Figure 3.**
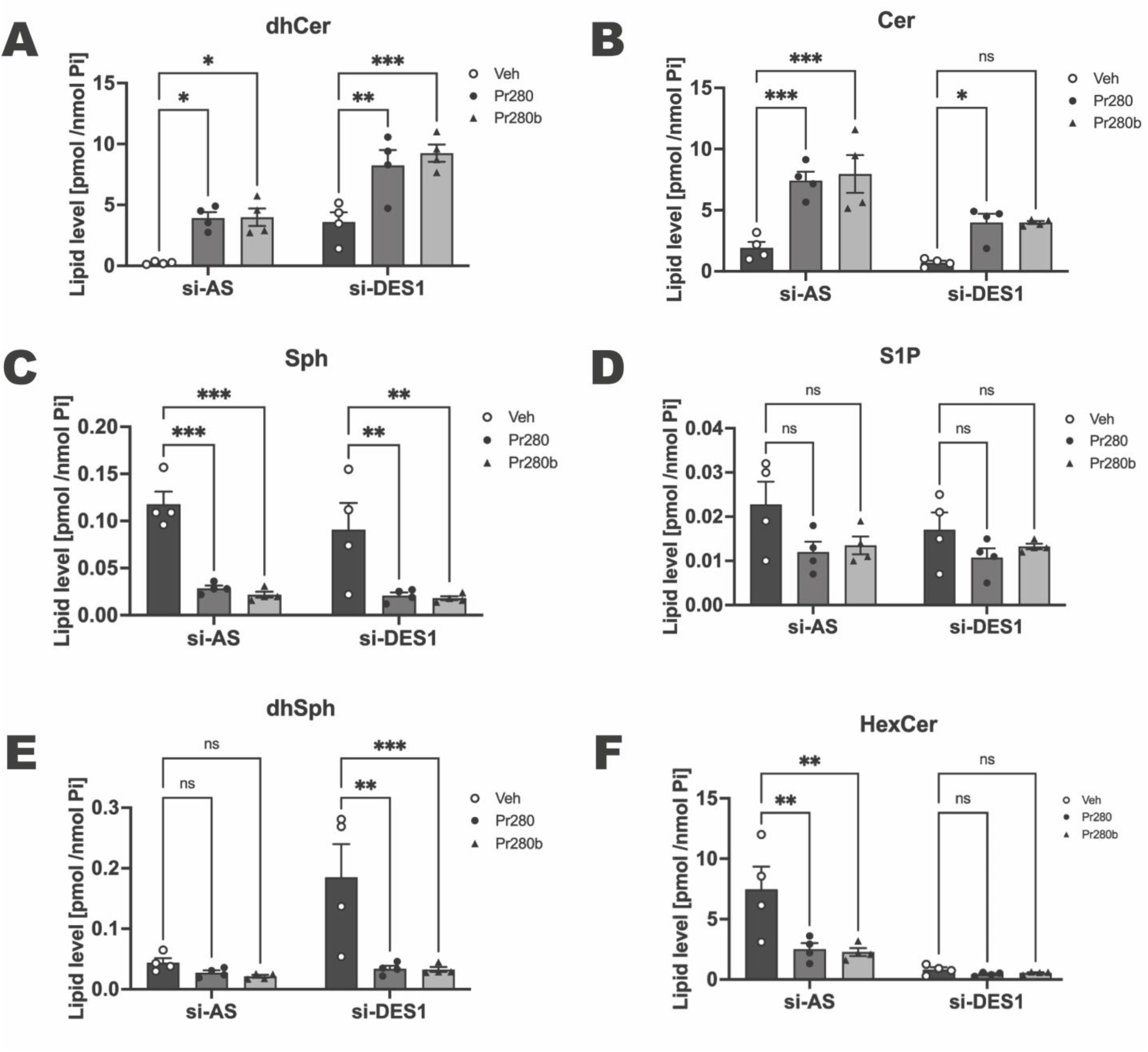
PR280 and PR280b have effects on lipids independently of DES1 status. JIMT1 cells were pre-treated for 48h with 20nM negative control (si-AS) or DES1 siRNA (si-DES1) prior to treatment with vehicle (ethanol), 2 μM PR280 or 2 μM PR280b for 24h. Lipids were extracted and analyzed for the lipid classes shown. Data are expressed as mean ± SEM of lipid level normalized to total inorganic lipid phosphate (n = 4; * p< 0.05; ** p < 0.01, *** p < 0.001 vs vehicle; two-way ANOVA with Dunnet’s post-test).

### PR280 and PR280b do not affect GCS or SM synthesis in the Golgi

Cer is the central hub of the sphingolipid network and serves as a precursor for multiple downstream sphingolipids, including sphingomyelin and hexosylceramides. Thus, we speculated that the unexpected increase in Cer observed with both compounds was due to inhibition of some of these metabolic pathways – particularly given that both compounds are Cer analogues. To investigate possible off-target effects, we utilized a recently developed NBD C6-Cer assay recently developed at Stony Brook. Here, the *in situ* conversion of NBD-C_6_-ceramide to NBD-C_6_-sphingomyelin and NBD-C_6_-hexosylceramides is measured by HPLC and, importantly, the assay allows the activities of Golgi-resident enzymes sphingomyelin synthases 1+2, ceramide kinase (CERK), and glucosylceramide synthase (GCS) to be measured simultaneously (35). For these studies, we utilized PR280 and PR280b at 2 μM and 5 μM doses for 24h as the time and doses used in the lipid experiments above **(Fig 4)**. In vehicle-treated cells, there was comparable flux through SMS (reflected in NBD-C_6_-sphingomyelin) and GCS (reflected in NBD-C_6_-hexosylceramides) with SMS activity being slightly higher, as seen in other cells (31). Importantly, no significant change in SM or GluCer synthesis was observed with either compound at either dose and, if anything, both compounds may have slightly elevated SM synthesis at higher doses. This suggests that neither compound is directly targeting the SMS or GCS activity under these experimental conditions and consequently, the increase in Cer is not through blocking of these pathways.

**Figure 4.**
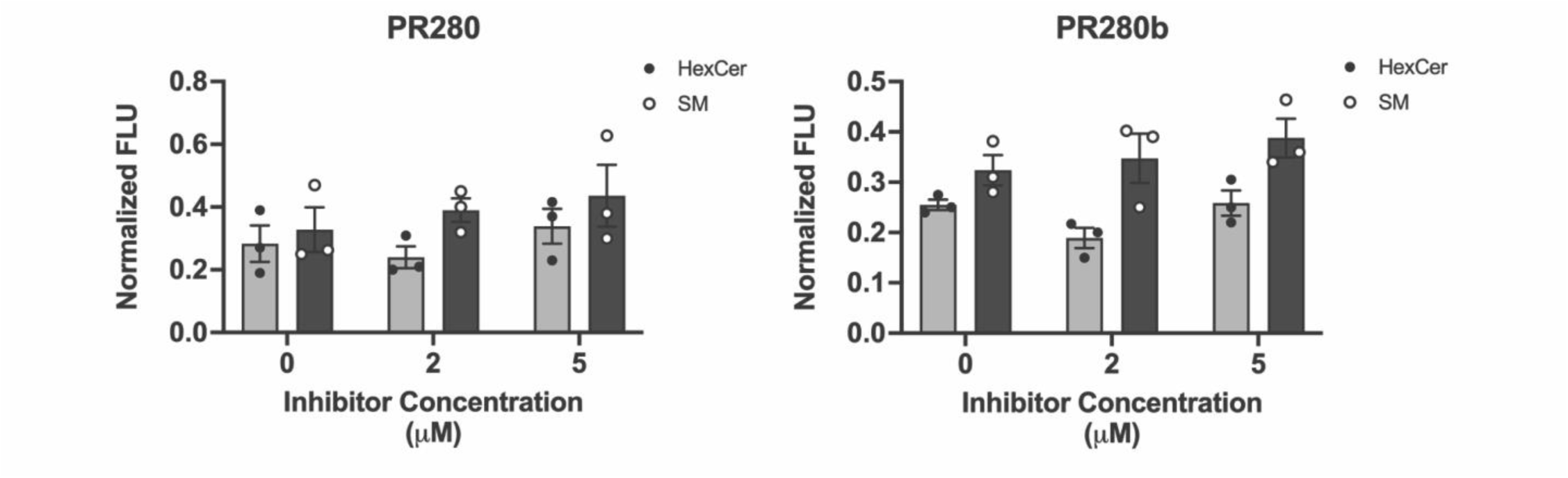
PR280 and PR280b do not inhibit Golgi SMS or GCS activity. JIMT1 cells were treated with vehicle (Ethanol), PR280, or PR280b for 24h with the doses shown. In situ SMS and GCS activity was analyzed using NBD-C6-Cer as described in Materials and Methods. Data are expressed as mean ± SEM normalized fluorescence units for NBD-HexCer and NBD-SM (n = 3)

### PR280 and PR280b modestly decreases AIS and CFA in breast cancer but do not affect other DES1-dependent biology

Despite the potential for off-target effects of these compounds, it was important to assess biological functions and specifically their effects on reported DES1-driven biologies. Previous studies from our lab showed that DES1 is crucial for anchorage-independent survival (AIS) – a key metastasis enabling biology – in HER2+ breast cancer cells and that targeting DES1 with either 4HPR – a previously reported DES1 inhibitor – or siRNA/Crispr knockout leads to a significant reduction in both AIS and soft agar colony formation assays (CFA) *in vitro* (14). For this, we utilized 2 μM of each compound as this had robust effects on DES activity and lower concentrations can reduce off-target effects. AIS assays showed that a modest – but statistically insignificant - increase in cell death in suspension with both compounds. However, there were no effects on viability in monolayer-plated cells **(Fig. 5A)** as seen before with DES1 siRNA and Crispr in HER2+ BC (14). To further investigate functional consequences, soft agar colony formation assays (CFA) were performed. These assays are a measure of a cancer cell’s tumorigenicity (aggressiveness) and test the cells’ ability to grow independently of an extracellular matrix, as cells are plated as singlets and proliferate into clusters after several weeks. As seen with trypan blue assays above, there was a modest effect of both compounds on both the number and size of colonies seen but these changes were not statistically significant **(Fig. 5B)**. Outside of AIS and colony formation, DES1 inhibition or loss has also been reported to have various signaling effects including apoptosis (7, 8) and induction of autophagy (11, 12). These effects have also been seen with other DES1 inhibitors such as 4HPR and XM462. Accordingly, we analyzed markers for apoptosis (PARP, Cleaved PARP) and autophagy (LC3) in cells plated in monolayer and treated with 2 μM of PR280 or PR280b for 24h (**Fig. 5C**). As can be seen, effects of either compound on PARP were modest with a modest decrease in total PARP levels compared to Vehicle and a slight increase in cleaved PARP levels. Analysis of LC3 suggested neither compound was able to stimulate appreciable LC3-II compared to vehicle control indicating that autophagy is not being induced. Taken together with the above results, these data suggest that at best, PR280 and PR280b are only modestly able to phenocopy biologies associated with loss of DES1 activity.

**Figure 5.**
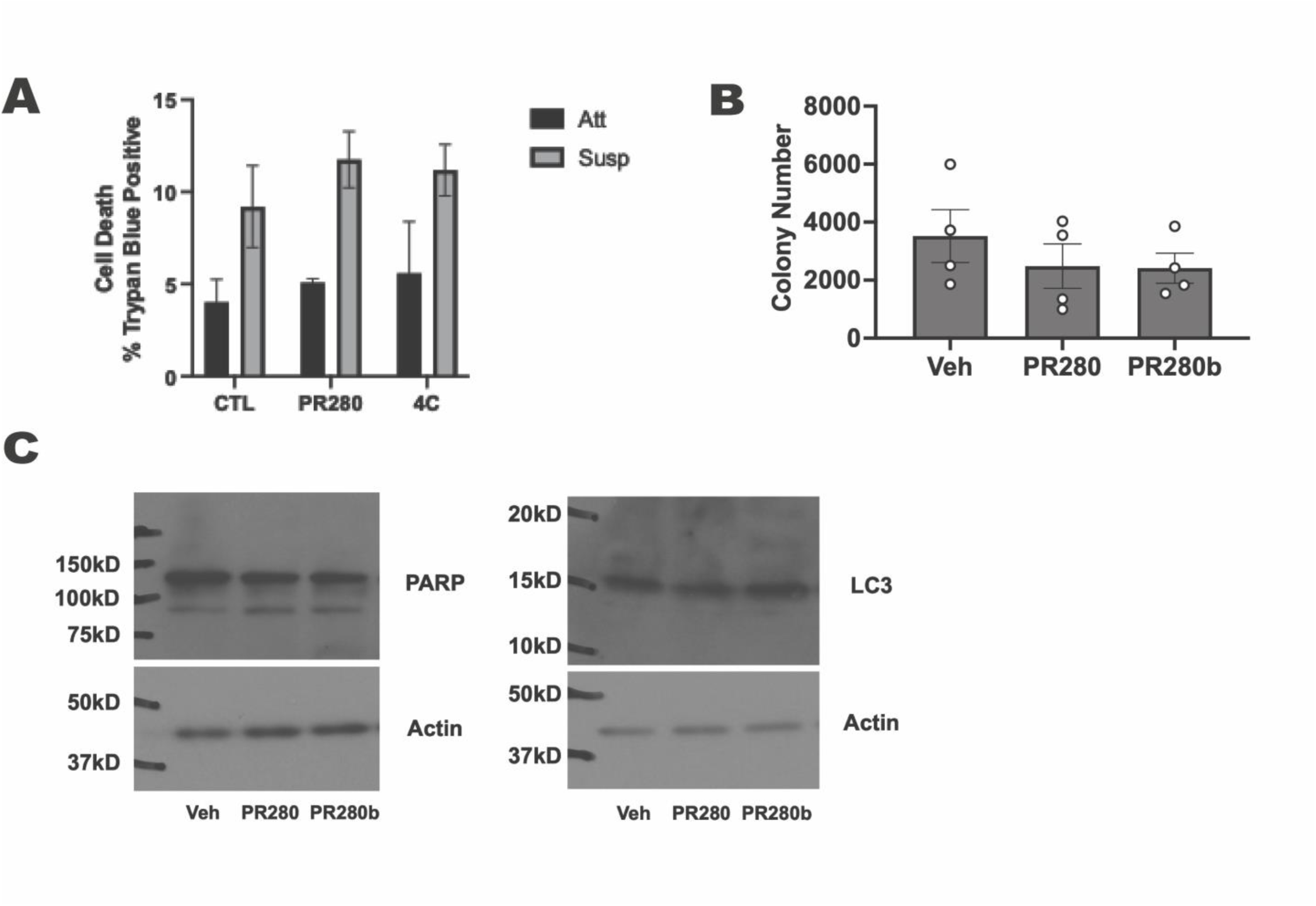
PR280 and PR280b have modest effects on reported DES1-mediated biologies and signaling. (A) JIMT1 cells were treated in monolayer (Att) or suspension (Susp) for 48h with vehicle, 2μM PR280 or 2μM PR280b. Cell viability was analyzed by trypan blue exclusion. Data are expressed as mean ± SEM % trypan blue positive (n = 3) (B) JIMT1 cells were plated in soft agar in the presence of vehicle, 2μM PR280 or 2μM PR280b. Colonies were cultured for 21-28d, stained, and quantified. Data are expressed as mean ± SEM % total colony count (n = 4) (C) JIMT1 cells in monolayer were treated with vehicle, 2mM PR280 or 2mM PR280b for 24h. Protein was extracted and analyzed by immunoblot as shown. Representative of at least 3 biological replicates

## DISCUSSION

The emerging roles of DES1 in various cellular biologies and diseases underscores the growing potential of DES1 inhibitors, both as research tools and therapeutics. In this study, we have further evaluated the efficacy, specificity, and biological activity of the recently reported DES1 inhibitor PR280 and its analogue PR280b. Notably, while both compounds are robust inhibitors of DES1 activity, they had broader effects on cellular SL metabolism and were unable to phenocopy reported biological effects associated with DES1 loss, suggesting off-target inhibition of other SL enzymes. Overall, while this limits the direct utility of PR280 as a specific DES1 inhibitor, these results highlight the potential of this scaffold as a promising basis for further inhibitor development – particularly given its inhibitory DES1 effects are greater than seen for existing DES1 inhibitors.

Despite growing evidence of its biological and pathological importance, DES1 has been of relatively limited interest as a therapeutic target compared to other SL enzymes e.g. sphingosine kinases as favored targets and effective DES1 inhibitors have been comparatively scarce and met with issues of stability or specificity (30, 32–34, 38). The cyclopropenone-containing Cer analog PR280 was recently developed as a novel DES1 inhibitor, based on structure-function studies of the prior inhibitors XM462 and GT11 (35). Here, we extend these findings to show that PR280 and its newly synthesized analogue PR280b are profound *in vitro* and *in situ* inhibitors of DES1 activity. Of note, our results show substantially stronger effects of parental PR280 than observed previously (35) showing >95% inhibition at 250nM *in vitro* in JIMT1 lysates compared to an IC50 of 700nM in T98 lysates. These differences may reflect the specific in vitro assay conditions being utilized or may be related to the cell lysate source being used. Nonetheless, the stronger inhibition seen here did also extend to in situ activity with both analogues showing >95% inhibition at 1μM – greater than the approximately 80% seen at 10 μM previously (35). Additionally, effects of both compounds on DES1 activity were rapid (<2h) and did not involve degradation of DES1 protein as has been reported previously for other DES1 inhibitory compounds (39). The inhibition of DES1 was further supported by lipidomic analysis showing robust dhCer accumulation following treatment and in line with DES1 KO and siRNA-treated cells. However, despite profound inhibition, effects on DES1-driven biologies and signaling were more mixed with both analogues showing only modest changes in AIS and CFA assays unlike what was seen previously with inhibitors and genetic DES1 loss (14). Similarly, neither compound had strong effects on apoptosis (PARP cleavage) or autophagy (LC3) despite both being known DES1-regulated biologies (7, 8) (11, 12). While this could be attributed to the specific cell type being studied, this more plausibly is due to off target effects. Indeed, broader analysis of SL levels clearly indicated some disconnect between inhibitor treated cells and genetic DES1 manipulation. In addition to increased Cer levels, both inhibitors had distinct effects on Sph and dhSph. Effects on SL beyond DES1 were further supported by experiments combining pharmacological treatment with DES1 knockdown with similar changes in Cer, Sph, and dhSph observed in the absence of DES1 protein. Collectively, this supports a mixed cellular response for both compounds including a potent DES1 inhibition but with additional broader effects on SL metabolism.

The off-target effects of PR280 and PR280b are not entirely unexpected given that they are Cer analogues. Indeed, the structural similarity to Cer suggests the strong likelihood they interact and modulate other SL enzymes as has been seen with other Cer analogues. For example, PDMP – originally developed as a GCS inhibitor – was reported to inhibit Cer glycanase (40), can inhibit Cer kinase (31) and induces lysosomal lipid accumulation, potentially beyond effects on GCS (41). Here, consideration of the sphingolipid-profiles in inhibitor treated cells with DES1 KO (**Fig. 2**) and siRNA treated cells (**Fig.3**) offers clues as to possible candidate off-target enzymes with the accumulation of Cer in both cases suggesting inhibition of a Cer-metabolizing enzyme. Here, our studies directly ruled out inhibition of SM synthase and GCS confirming that the observed decreases in HexCer (**Fig. 2F**) and SM (**Fig. 2G**) are likely consequences of reduced downstream flux, supported by the similar effects of DES1 KO. Similarly, comparable effects of inhibitors and DES1 KO/siRNA on S1P suggest that neither analog is impacting the sphingosine kinases (SK1/2) – a valid concern given the inhibition of DES1 seen with SKi-II and the SK2 inhibitor ABC294640 (39, 42). Conversely, as both inhibitors significantly reduced sphingosine levels, more so than both DES1 siRNA or Crispr-mediated KO, this implies potential inhibition of ceramidase enzymes. Indeed, many reported ceramidase inhibitors are Cer analogues including D-MAPP (43), LCL521 (44), Urea-Cer (45), and KPB82 (46). However, we also cannot rule out the possibility that altered synthesis or recycling through Cer synthases could also contribute – particularly given both compounds did not lead to accumulation of dhSph, unlike genetic DES1 manipulations. Given that both compounds may influence several connected aspects of SL metabolism, measurements of steady-state lipid abundance may not be sufficient to identify the responsible reactions or establish changes in metabolic flux. Thus, a more systematic approach may be needed to define the key off-target enzymes.

In conclusion, PR280 and PR280b are promising inhibitors that achieve substantial DES1 inhibition at low concentrations, but with cellular effects extend beyond the lipid consequences of genetic DES1 depletion alone. Defining compound metabolism, measuring SL flux, and resolving changes in individual lipid species will help distinguish DES1-dependent effects and help identify the key off-target enzymes of both compounds. This would allow for future structure-function analysis that could guide the development of analogues that maintain strong inhibitory potency of DES1 with increased specificity. Given the growing evidence linking DES1 to various diseases, such inhibitors would be particularly valuable as candidate therapeutics.

## DATA AVAILABILITY

Requests for further information on reagents, methodologies and data should be directed to Christopher Clarke

## ACKNOWLEDGEMENTS

The authors wish to acknowledge the Stony Brook Cancer Center Biological Mass Spectrometry Shared Resource and its staff and personnel for expert assistance with lipid measurements. Funding for this project came from National Institutes of Health grant R01 CA248014. We also thank Grant PID2021-125923OB-I00 funded by MCIN/AEI/10.13039/501100011033 and by “ERDF A way of making Europe”. P.R. acknowledges Universitat Rovira i Virgili for a predoctoral fellowship (2019PMF-PIPF-36), as well as Secretaria d’Universitats i Recerca de la Generalitat de Catalunya and ESF for a predoctoral FI fellowship (FI2021/B00241).

## AUTHOR CONTRIBUTIONS

C.J.C. and M.I.M conceptualized and supervised the project. D.M.P., H.R.R.G., C.N.C., and N.M. performed the experiments and analyzed the data. P.R., Y.D., and M.I.M. synthesized PR280 and its analogues. D.M.P., M.I.M., and C.J.C. wrote the original draft and made figures. All authors read the draft and provided input.

