## Supplemental Information for "Evaluation of the Biological Effects and Specificity of the Novel Dihydroceramide Desaturase 1 inhibitor PR280 and its analogues"


#### Table of Contents


1. General synthetic methods........................................................................S2
2. Synthetic procedures ..............................................................................S3
3. $^{1}H$ and $^{13}C$ NMR spectra ..................................................................S5

1. **General synthetic methods**

All reactions were carried out in anhydrous conditions using standard vacuum-argon techniques. Reaction flasks were flame-dried under vacuum using a heat gun and then purged with argon. Reagents and solvents were transferred using cannulas or syringes previously flushed with argon.

IR spectra were recorded on a JASCO FT/IR-680 Plus Fourier-transform infrared spectrophotometer and processed using SpectraManager software (JASCO®). Optical rotations ($[\alpha]_D^{25}$) were measured on a Perkin-Elmer 241 polarimeter using a 1.0 dm-path length cell. Melting points (m.p.) were determined on a Mettler Toledo DSC 822e instrument. High–resolution mass spectra (HRMS) were recorded on an Agilent 1100 Series LC/MSD mass spectrometer equipped with an electrospray ionization (ESI) source; exact *m*/*z* values are reported in Daltons (error < 5 ppm). $^{1}$H, and $^{13}$C NMR spectra were acquired on a Varian Mercury VX 400 spectrometer and processed using MestreNova software (MestreLab®). Coupling constants (*J*) are reported in Hz. Signal multiplicities are denoted as follows: s = singlet, d = doublet, t = triplet, q = quartet, p = quintet, m = multiplet, dd = doublet of doublets, ap = apparent, b = broad. Chemical shifts are referenced to the residual solvent signals. Thin-layer chromatography (TLC) was performed on E. Merck® aluminum plates (0.25 mm) coated with silica gel 60 F254, with visualization by *p*-anisaldehyde or $KMnO_4$ staining as indicated. Flash column chromatography was carried out using Fluka® or Merck® silica gel 60 (230–400 mesh), eluting with the specified solvents under pressure.

**2. Synthetic procedures**

Cyclopropenone ceramide analogues **PR280a** and **PR280b** were synthesized as described for the reference compound **PR280** (ref. 34 in the main article). Additional information is provided below.

***N*-((1*R*,2*S*)-1,3-dihydroxy-1-(3-oxo-2-tridecylcycloprop-1-en-1-yl)propan-2-yl)hexanamide (PR280a)**

Hexanoic acid (21.5 µL, 0.17 mmol), EDC (33.2 mg, 0.17 mmol) and HOBt (23.3 mg, 0.17 mmol) were dissolved in dry $CH_2Cl_2$ (4.9 mL). The solution was stirred at room temperature for 45 min. The resulting mixture was added to a solution of aminodiol **1** (50.9 mg, 0.16 mmol) in dry $CH_2Cl_2$ (5.4 mL) and stirred at room temperature overnight.

Upon completion of the reaction, water was added, and the phases were separated. The aqueous phase was extracted with $CH_2Cl_2$, and the combined organic extracts were dried over anhydrous $Na_2SO_4$, filtered, and concentrated under reduced pressure to furnish a yellowish solid residue. Purification by flash column chromatography on silica gel ($CH_2Cl_2$/MeOH/$NH_4OH$, 96:4:1), followed by recrystallization from pentane afforded the desired ceramide analogue **PR280a** as a white solid (48.6 mg, 0.11 mmol, 73 % yield).

***R*f** = 0.37 ($CH_2Cl_2$/MeOH/$NH_4OH$ (9:1:0.1)). $[\alpha]_D^{25}$-33.9 (c 0.6, *$CHCl_3$*); **m.p.** = 121-122 °C; **$^1$H NMR** δ in ppm (400 MHz, $CDCl_3$): 6.97 (d, *J* = 5.9 Hz, 1H), 5.94 (d, *J* = 7.5 Hz, 1H), 4.85-4.83 (m, 1H), 4.26-4.23 (m, 1H), 4.22-4.17 (m, 1H), 3.99-3.93 (m, 1H), 3.90-3.84 (m, 1H) 2.66 (t, *J* = 7.4 Hz, 2H), 2.28-2.16 (m, 2H), 1.71 (p ap, *J* = 7.2 Hz, 2H), 1.59 (p ap, *J* = 7.3 Hz, 2H), 1.38-1.23 (m, 24 H), 0.88 (t, *J* = 7.0 Hz, 3H), 0.88 (t, *J* = 7.0 Hz, 3H); **$^{13}$C NMR** δ in ppm (100.6 MHz, $CDCl_3$): 176.5, 162.2, 160.6, 159.5, 72.1, 61.5, 56.5, 36.3, 32.1, 32.0, 29.82, 29.78, 29.76, 29.6, 29.5, 29.4, 29.3, 26.6, 26.2, 25.4, 22.8, 22.5,

14.3, 14.0; **HRMS** (ESI+) for $[M+H]^+$ $C_{25}H_{46}NO_4^+$ (m/z): calculated: 424.34214, found: 424.34103; **FT-IR (ATR)** ν in $cm^{-1}$: 3303, 2953, 2851, 1838, 1645, 1577, 1547, 1466, 1457, 1417, 1376, 1326, 1289, 1254, 1217, 1120, 1071, 980, 952, 720, 695, 652, 620, 437.

**_N_-((1_R_,2_S_)-1,3-dihydroxy-1-(3-oxo-2-tridecylcycloprop-1-en-1-yl)propan-2-yl)decanamide (PR280b)**

Decanoic acid (29.3 mg, 0.17 mmol), EDC (32.7 mg, 0.17 mmol) and HOBt (23.2 mg, 0.17 mmol) were dissolved in dry $CH_2Cl_2$ (4.9 mL). The solution was stirred at room temperature for 45 min. The resulting mixture was added to a solution of aminodiol **1** (50.9 mg, 0.16 mmol) in dry $CH_2Cl_2$ (5.4 mL) and stirred at room temperature overnight.

Upon completion of the reaction, water was added, and the phases were separated. The aqueous phase was extracted with $CH_2Cl_2$, and the combined organic extracts were dried over anhydrous $Na_2SO_4$, filtered, and concentrated under reduced pressure to furnish a yellowish solid residue. Purification by flash column chromatography on silica gel ($CH_2Cl_2$/MeOH/$NH_4OH$, 96:4:1), followed by recrystallization from pentane afforded the desired ceramide analogue **PR280b** as a white solid (49.6 mg, 0.10 mmol, 66 % yield).

***R*f** = 0.40 ($CH_2Cl_2$/MeOH/$NH_4OH$ (9:1:0.1)). $[\alpha]_D^{25}$-10.6 (c 0.4, *$CHCl_3$/MeOH*, 1:1); **m.p.** = 132-133 °C; **$^1$H NMR** δ in ppm (400 MHz, $CDCl_3$): 6.79 (d, *J* = 5.6 Hz, 1H), 5.84 (bs, 1H), 4.82 (bs, 1H), 4.22-4.18 (m, 1H), 4.00 (dd, *J* = 11.3 Hz, *J* = 4.3 Hz, 1H), 3.90 (dd, *J* = 11.3 Hz, *J* = 5.0 Hz, 1H), 2.67 (t, *J* = 7.5 Hz, 2H), 2.29-2.17 (m, 2H), 1.75-1.68 (m, 2H), 1.59 (p ap, *J* = 7.3 Hz, 2H), 1.38-1.26 (m, 32H), 0.88 (t, *J* = 6.5 Hz, 6H); **$^{13}$C NMR** δ in ppm (100.6 MHz, $CDCl_3$): 176.8, 162.2, 160.6, 159.4, 72.8, 61.5, 56.4, 36.4, 32.1, 32.0, 29.83, 29.80, 29.78, 29.62, 29.60, 29.51, 29.49, 29.43, 29.36, 29.3, 26.6, 26.3, 25.79, 22.84, 22.81, 14.3, 14.2; **HRMS** (ESI+) for $[M+H]^+$ $C_{29}H_{54}NO_4^+$ (m/z): calculated: 480.40474, found: 480.40344; **FT-IR (ATR)** ν in $cm^{-1}$: 3306, 2955, 2919, 2851, 1838, 1647, 1577, 1547, 1467, 1417, 1389, 1330, 1292, 1259, 1230, 1200, 1117, 1073, 981, 944, 721, 695, 654, 621, 537.

### 3. $^{1}$H and $^{13}$C NMR spectra

***N*-((1*R*,2*S*)-1,3-dihydroxy-1-(3-oxo-2-tridecylcycloprop-1-en-1-yl)propan-2-yl)hexanamide (PR280a)**

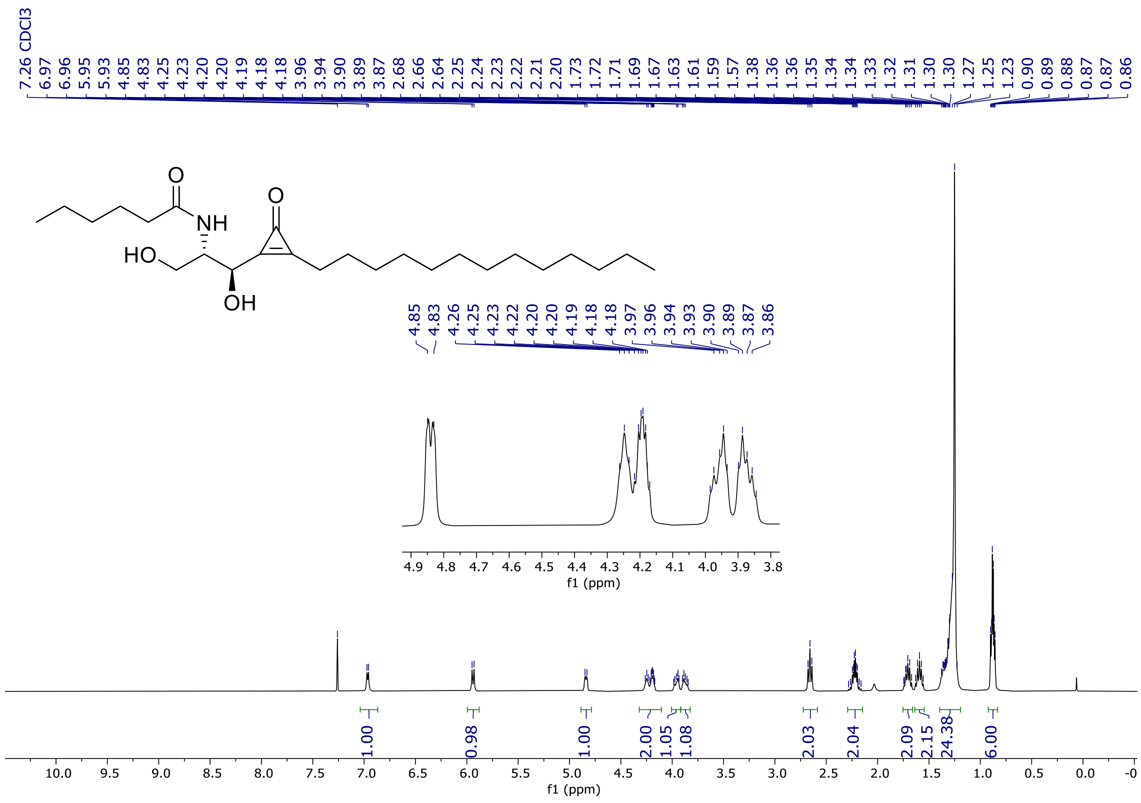


**Figure S1.** $^{1}$H NMR (400 MHz, $CDCl_3$) of **PR280a**.

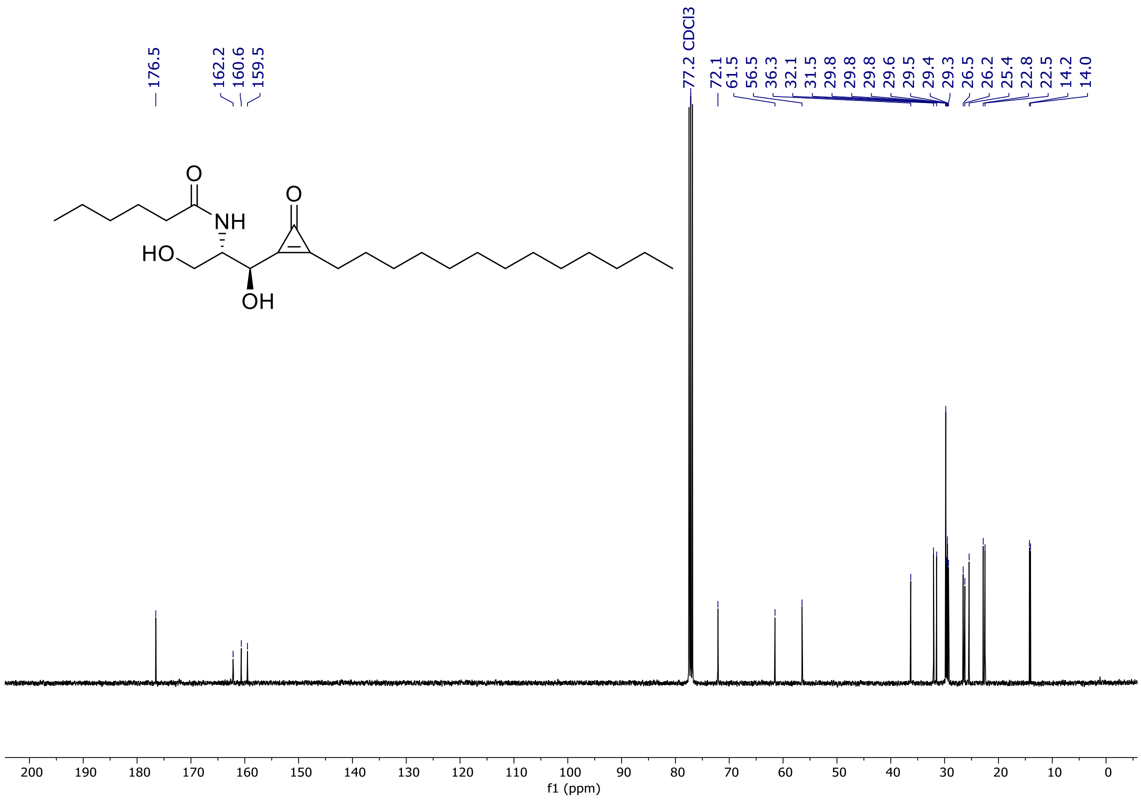


**Figure S2.** $^{13}$C NMR (100.6 MHz, $CDCl_3$) of **PR280a**.

***N*-((1*R*,2*S*)-1,3-dihydroxy-1-(3-oxo-2-tridecylcycloprop-1-en-1-yl)propan-2-yl)decanamide (PR280b)**.

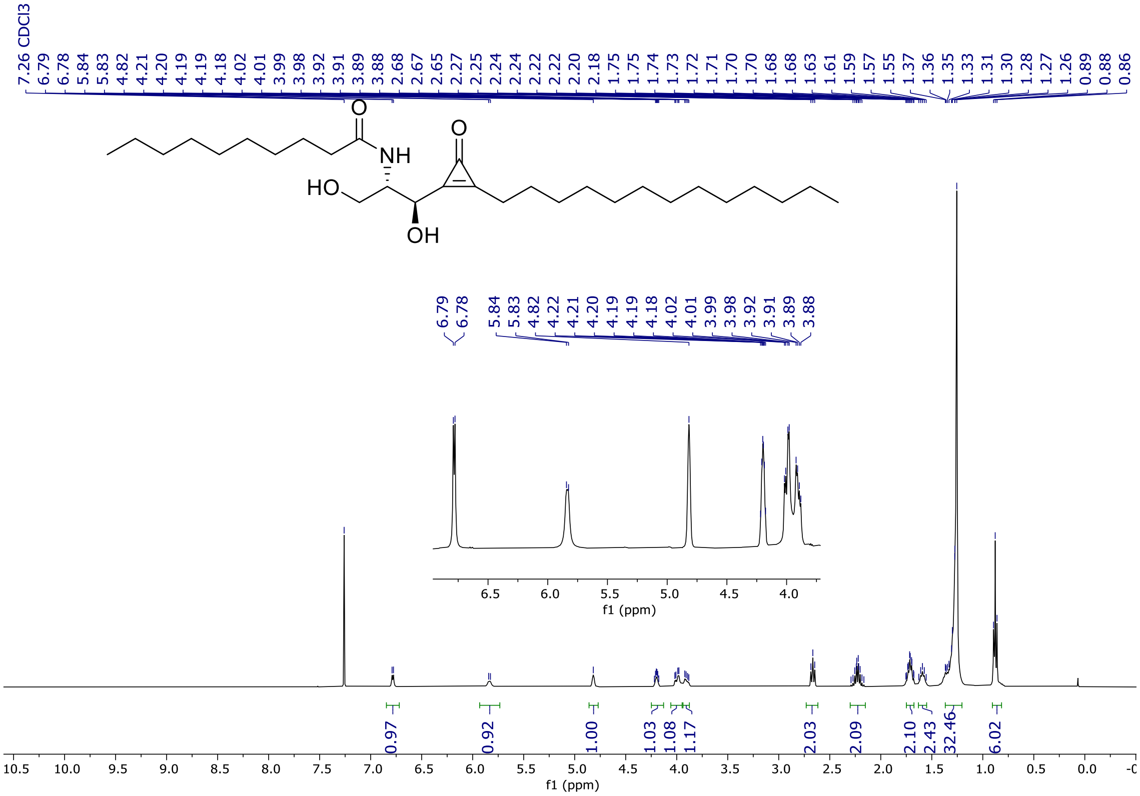


**Figure S3.** **$^1$H NMR** (400 MHz, $CDCl_3$) of **PR280b**.

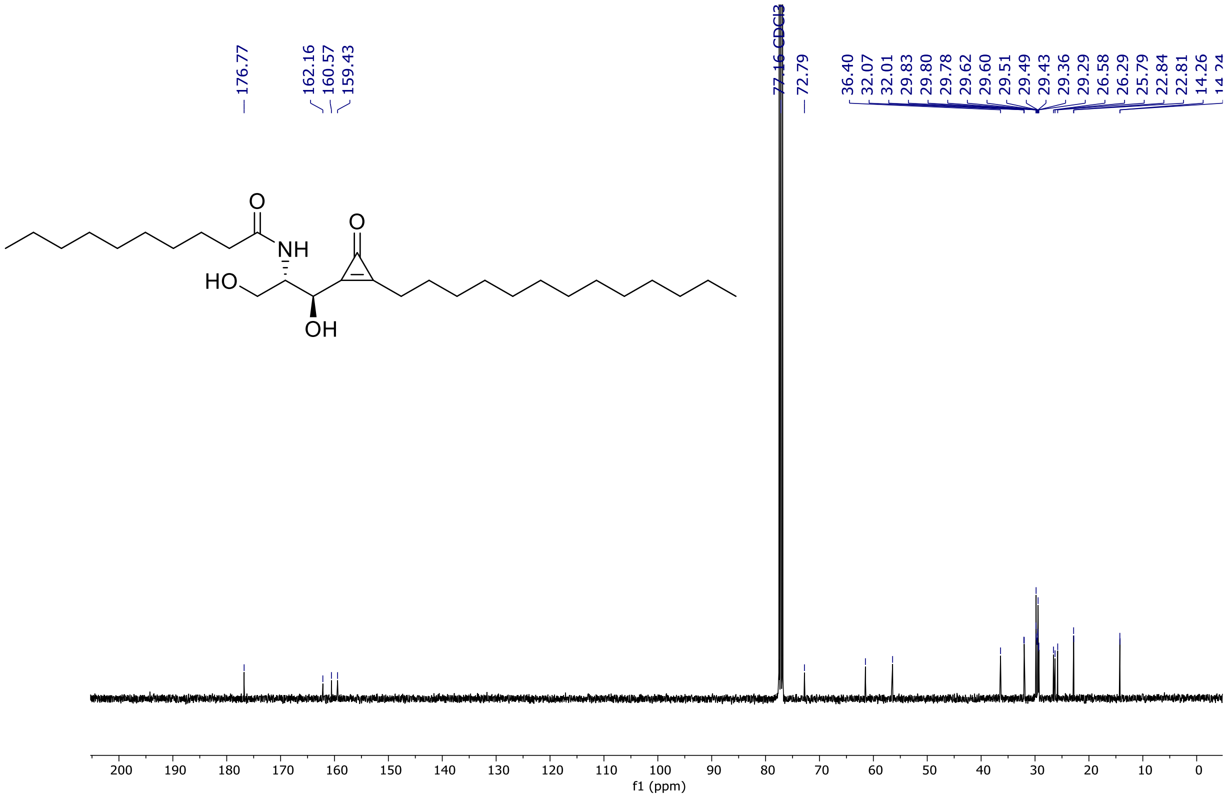


**Figure S4.** $^{13}$C NMR (100.6 MHz, $CDCl_3$) of **PR280b**.
